# Long solids retention times and attached growth phase favor prevalence of comammox bacteria in nitrogen removal systems

**DOI:** 10.1101/696351

**Authors:** Irmarie Cotto, Zihan Dai, Linxuan Huo, Christopher L. Anderson, Katherine J. Vilardi, Umer Ijaz, Wendell Khunjar, Christopher Wilson, Haydee De Clippeleir, Kevin Gilmore, Erika Bailey, Ameet J. Pinto

## Abstract

The discovery of the complete ammonia oxidizing (comammox) bacteria overturns the traditional two-organism nitrification paradigm which largely underpins the design and operation of nitrogen removal during wastewater treatment. Quantifying the abundance, diversity, and activity of comammox bacteria in wastewater treatment systems is important for ensuring a clear understanding of the nitrogen biotransformations responsible for ammonia removal. To this end, we conducted a yearlong survey of 14 full-scale nitrogen removal systems including mainstream conventional and simultaneous nitrification-denitrification and side-stream partial nitrification-anammox systems with varying process configurations. Metagenomics and genome-resolved metagenomics identified comammox bacteria in mainstream conventional and simultaneous nitrification-denitrification systems, with no evidence for their presence in side-stream partial nitrification-anammox systems. Further, comammox bacterial diversity was restricted to clade A and these clade A comammox bacteria were detected in systems with long solids retention times (>10 days) and/or in the attached growth phase. Using a newly designed qPCR assay targeting the *amoB* gene of clade A comammox bacteria in combination with quantitation of other canonical nitrifiers, we show that long solids retention time is the key process parameter associated with the prevalence and abundance of comammox bacteria. The increase in comammox bacterial abundance was not associated with concomitant decrease in the abundance of canonical nitrifiers; however, systems with comammox bacteria showed significantly better and temporally stable ammonia removal compared to systems where they were not detected. Finally, in contrast to recent studies, we do not find any significant association of comammox bacterial prevalence and abundance with dissolved oxygen concentrations in this study.

**Highlights:**

- Clade A comammox bacteria were detected in wastewater nitrogen removal systems.
- New qPCR assay targeting the *amoB* gene of clade A comammox bacteria was developed.
- Comammox bacteria are prevalent in mainstream conventional and simultaneous nitrification-denitrification systems with long solids retention times (>10 days).
- Comammox bacteria were not detected in sidestream partial nitrification-anammox systems included in this study.

GRAPHICAL ABSTRACT

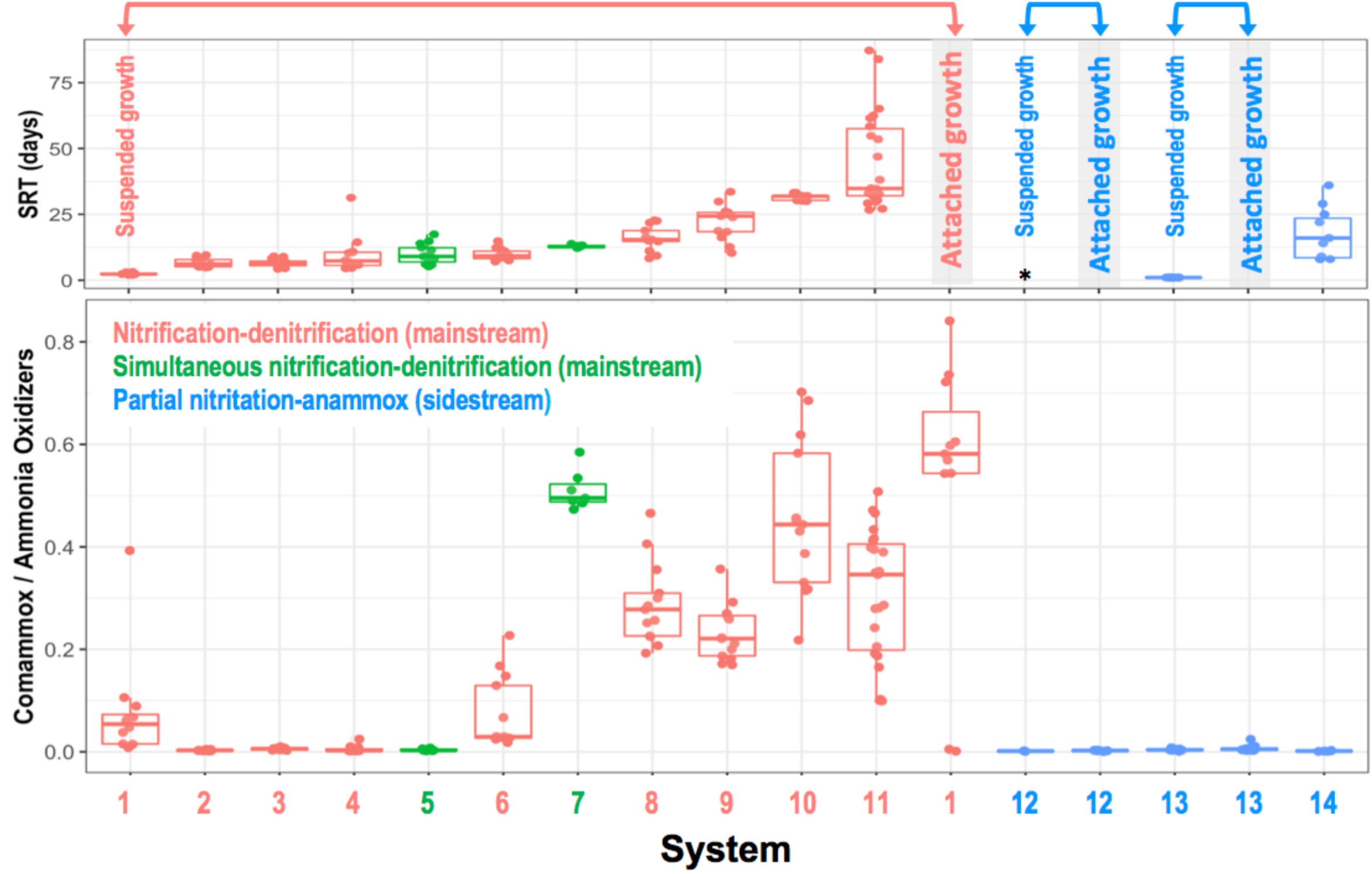

## 1.0 Introduction

Nitrification, the oxidation of ammonia to nitrate via nitrite, coupled with denitrification where nitrate is reduced to dinitrogen gas are key processes in the removal of nitrogen from wastewater (Klotz and Stein 2008). Traditionally, nitrification was considered as a two-step process driven by two distinct nitrifying guilds, i.e., ammonia oxidation by the aerobic ammonia oxidizing bacteria (AOB) (Kowalchuk and Stephen 2001) or archaea (AOA) (Stahl et al. 2012) followed by nitrite oxidation by the aerobic nitrite oxidizing bacteria (NOB) ((Daims et al. 2016). Ammonia oxidizers within the *Betaproteobacteriales* genera *Nitrosomonas* and *Nitrosospira* (Siripong and Rittmann 2007, Wu et al. 2019) and NOB within the genus *Nitrospira* (Juretschko et al. 1998, Wu et al. 2019) and more recently *Nitrotoga* (Saunders et al. 2016) are thought to be dominant nitrifiers in wastewater treatment systems. However, the discovery of complete ammonia oxidizing (i.e., comammox) bacteria (Daims et al. 2015, Pinto et al. 2015, van Kessel et al. 2015) has added new complexity to nitrogen biotransformation in wastewater systems. For instance, comammox bacteria can completely oxidize ammonia to nitrate and may compete with canonical AOB and NOB, as well as anaerobic ammonia oxidizing (anammox) bacteria in partial nitritation-anammox (PNA) systems, also known as deammonification systems. The potential competition for ammonia and possibly nitrite among comammox bacteria and other nitrifiers could have implications for process design and operation not only in wastewater treatment but also across other ecosystems.

Comammox bacteria have been detected in geothermal springs (Daims et al. 2015), aquaculture systems (van Kessel et al. 2015), drinking water (Pinto et al. 2015, Tatari et al. 2017, Wang et al. 2017), rapid gravity sand filters treating groundwater (Fowler et al. 2018, Palomo et al. 2016), soils (Hu and He 2017, Orellana et al. 2018), as well as a range of wastewater treatment bioreactors (Annavajhala et al. 2018, Camejo et al. 2017, Chao et al. 2016, Fan et al. 2017, Gonzalez-Martinez et al. 2016, Pjevac et al. 2017, Roots et al. 2019, Spasov et al. 2019, Wang et al. 2018, Xia et al. 2018). The key to evaluating the impact of comammox bacteria on wastewater process operations is to understand the impact of key process variables on whether comammox prevalence and abundance and in turn how this impacts overall activity and function of the engineered system. This can then help delineate laboratory-, pilot-, or even full-scale experiments to probe competitive dynamics between comammox bacteria and other nitrifiers in scenarios that are relevant from a process operations perspective. To address this issue, the current study presents a systematic year-long evaluation of nitrifying populations, including comammox bacteria, in full-scale wastewater treatment plants to provide a baseline of process configurations, operations, and environmental conditions under which comammox bacteria might be important.

All detected comammox bacteria belong to genus *Nitrospira* (lineage II) and exhibit close phylogenetic relatedness to canonical *Nitrospira*-NOB. Recent work suggests that the capacity to oxidize ammonia via the ammonia monoxygenase (AMO) and hydroxylamine dehydrogenase (HAO) enzymes may have been acquired by comammox *Nitrospira* via horizontal gene transfer (Palomo et al. 2018). Nonetheless, the close phylogenetic affiliation of comammox bacteria with canonical NOB represents a major challenge with their detection and quantitation. Specifically, the 16S rRNA gene and subunits A (*nxrA*) and B (*nxrB*) of the nitrite oxidoreductase (NXR) genes cannot distinguish between comammox bacteria from canonical NOB within the genus *Nitrospira*. One alternative to identify and obtain relative abundance of comammox bacteria within a complex nitrifying consortium involves the use of shotgun DNA sequencing (i.e., metagenomics). This has been employed by several studies (Annavajhala et al. 2018, Camejo et al. 2017, Chao et al. 2016, Fan et al. 2017, Gonzalez-Martinez et al. 2016, Pinto et al. 2015, Roots et al. 2019, Spasov et al. 2019, Xia et al. 2018) to demonstrate that comammox bacterial presence in wastewater systems is primarily dominated by clade A comammox bacteria closely related to *Ca* Nitrospira nitrosa. While metagenomics provides a snapshot overview, it is not ideally suited for high-throughput profiling of large number of samples particularly for microbial groups that are typically low-to-medium abundance (e.g., nitrifiers) due to sequencing cost and sequencing depth issues. To this end, the ammonia monooxygenase (amo) gene sequences provide a convenient approach for detection and quantitation. Specifically, both the subunit’s A (*amoA*) and B (*amoB*) of the ammonia monooxygenase genes form distinct clusters from other known AOB and AOA. This sequence divergence has been used to develop primer sets targeting the *amoA* gene of clade A and clade B comammox bacteria separately (Pjevac et al. 2017) and together (Bartelme et al. 2017, Fowler et al. 2018, Wang et al. 2018). A consistent challenge with these primers is the formation of unspecific products and our experience in the study suggests that they are also unable to capture comammox bacteria detected via metagenomics (see results section). Alternatively, (Beach and Noguera 2019) proposed the use of species specific primers that target comammox bacterial *amoA* gene depending on the process of interest. For instance, they proposed that since *Ca* Nitrospira nitrosa are dominant in low energy wastewater treatment systems, utilizing primers that capture *Ca* Nitrospira nitrosa would be ideal. While this circumvents the challenge of unspecific product formation, a species-specific approach eliminates the possibility of detecting other closely related comammox bacteria. In this study, we used a metagenomic approach to recover *amoA* and *amoB* genes from several full-scale nitrogen removal systems and use them in combination with previously published gene sequences to design and validate clade level primers for the *amoB* gene of clade A comammox bacteria; this is to our knowledge the primary comammox clade of relevance of wastewater treatment systems.

The overall objectives of this study were (1) to identify nitrogen removal process configurations for wastewater treatment where comammox bacteria are likely to be relevant, (2) develop a qPCR assay for quantitation of comammox bacteria in complex nitrifying communities, and (3) perform a temporal survey across a range of process configurations to determine the influence of environmental and variable process operation conditions on the abundance of nitrifying populations inclusive of comammox bacteria. This was accomplished through year-long quantitative tracking of nitrifying populations in fourteen nitrogen removal systems with varying process configuration using genome resolved metagenomics and qPCR combined with appropriate statistical analyses.

## 2.0 Materials and Methods

### 2.1 Sampling sites, sample processing, and data collection

Samples were collected on a monthly basis from June 2017 to June 2018 at fourteen nitrogen removal systems which included seven full-scale suspended growth nitrification-denitrification (ND) systems, one full-scale integrated fixed film activated sludge (IFAS) ND system, two full-scale simultaneous ND (SND) systems, one full-scale suspended growth side stream PNA system, and two full-scale IFAS PNA systems. Table 1 provides an overview of the nitrogen removal systems sampled in this study. Sample collection and processing were conducted according to the MIDAS field guide (McIlroy et al. 2015). Specifically, samples were collected from the nitrifying bioreactors by operational personnel at each of the wastewater utilities and 100 ml of sample was shipped overnight in coolers with icepacks to Northeastern University (NU). Immediately upon arrival to NU, the samples were homogenized using the Hei-TORQUE Value 400 (Heidolph, Cat. No. 036093070) in a 30-mL glass/Teflon tissue grinder (DWK Life Sciences Wheaton, Cat. No. 357984) for 1 minute (2nd gear, speed 9, 10 times from top to bottom of the glass tissue grinder) following the MIDAS protocol, and four 1 mL homogenized aliquots per sample were transferred to Lysing Matrix E tubes (MP Biomedical, Cat. No. 6914100). Samples were centrifuged at 10,000g for 5 minutes and the supernatant was discarded and biomass pellet was stored at −80°C until DNA extraction. Further, ∼20 mg of media attached biomass samples from the IFAS system were also stored in Lysing Matrix E tubes at −80°C. In addition, the utilities provided data on pH, temperature, ammonia, nitrite, nitrate, chemical oxygen demand (COD), biological oxygen demand (BOD), total suspended solids (TSS), volatile suspended solids (VSS), mixed liquor TSS (MLSS) and mixed liquor VSS (MLVSS) from the influent, nitrification reactor, and effluent for each sampling time point, as well as hydraulic retention time (HRT), solids retention time (SRT) and dissolved oxygen (DO) data from the nitrification reactors.

**Table 1:** Overview of sampling locations, sampling scheme used in this manuscript, process type and sub-type, operational scale and sampling time-frame for this study.

| Site code | Process type | Process sub-type | Treatment stream | Sampling Start (mm/yy) | Sampling End (mm/yy) |
| --- | --- | --- | --- | --- | --- |
| NEU | ND<sup>1</sup> | Four/five stage | Mainstream | 06/17 | 06/18 |
| DUR | PNA<sup>2</sup> | Anitamox | Sidestream | 06/17 | 06/18 |
| YAN | SND<sup>3</sup> | ORP control<sup>a</sup> | Mainstream | 11/17 | 05/18 |
| GRE | ND<sup>1</sup> | SBR<sup>b</sup> | Mainstream | 06/17 | 06/18 |
| DCWM | ND<sup>1</sup> | Five stage | Mainstream | 06/17 | 06/18 |
| KIN | ND<sup>1</sup> | MBR/MLE<sup>c</sup> | Mainstream | 06/17 | 06/18 |
| WIL | ND<sup>1</sup> | Five stage | Mainstream | 06/17 | 06/18 |
| JAMM | ND<sup>1</sup> | IFAS<sup>d</sup> | Mainstream | 06/17 | 06/18 |
| JAMS | PNA<sup>2</sup> | Anitamox | Sidestream | 06/17 | 06/18 |
| YORM | ND<sup>1</sup> | Conventional | Mainstream | 06/17 | 06/18 |
| YORS | PNA<sup>2</sup> | DEMON | Sidestream | 06/17 | 06/18 |
| BOA | ND<sup>1</sup> | High MLSS | Mainstream | 06/17 | 06/18 |
| NAN | SND<sup>3</sup> | Five stage | Mainstream | 06/17 | 06/18 |
| ARM | ND<sup>1</sup> | Five stage | Mainstream | 06/17 | 06/18 |

<sup>1</sup>Nitrification-denitrification, <sup>2</sup>Partial nitrification-anammox, <sup>3</sup>Simultaneous nitrification-denitrification, <sup>a</sup>Oxidative-Reduction Potential Control, <sup>b</sup>Sequencing batch reactor, <sup>c</sup>Membrane bioreactor/Modified Ludzack-Ettinger, <sup>d</sup>Integrated Fixed-film Activated Sludge

### 2.2 DNA extraction

Samples were subject to DNA extraction using the DNAeasy PowerSoil kit (Qiagen, Inc., Cat. No.12888) with automated extraction instrument QIAcube (Qiagen, Inc., Cat. No. 9002160) following manufacturer instructions with some modifications in the sample preparation step. Briefly, the lysing buffer from the PowerBead Tubes provided in the DNAeasy PowerSoil kit was transferred to the lysing matrix tubes containing the samples and 60 µL of Solution C1 from the kit was added to each sample. To complete cell lysis, bead-beating was performed using a FastPrep-24 instrument (MP Bio, Cat. No. 116005500) four times for 40 seconds each (McIlroy et al. 2015). Between each 40 second bead beating interval, samples were kept on ice for 2 min to prevent excess heating. Samples were centrifuged at 10,000g for 30 seconds and the supernatant was further purified using the QIAcube Protocol Sheet for the DNeasy PowerSoil Kit. Ten samples and two blanks (only reagents) were extracted for each DNA extraction run. Extracted DNA was quantified using Qubit instrument with Qubit dsDNA Broad Range Assay (ThermoFisher Scientific, Cat. No. Q32850) and a subset of samples were randomly selected for analysis using 1% agarose gel electrophoresis to visualize DNA shearing. Extracted DNA concentrations ranged from 2 to 254 ng/µL. DNA extracts were stored at −80°C until further analysis.

### 2.3 Metagenomic sequencing and data analyses

One sample from each system was selected for metagenomic sequencing based on qPCR estimates of AOB and *Nitrospira* (see qPCR details in section 2.4). Specifically, samples with high *Nitrospira*:AOB ratios were selected for metagenomic sequencing under the assumption that these samples were likely to consist of comammox *Nitrospira* bacteria. For systems that included distinct suspended and attached growth phase (e.g., IFAS system from JAMM and PNA systems from JAMS and DUR), one sample each for the attached and suspended phase were included in the sequencing run resulting in a total of 18 metagenomes. Extracted DNA was shipped frozen to the Roy J. Carver Biotechnology Center at University of Illinois Urbana-Champaign Sequencing Core for sequencing. The DNA extracts were subject to PCR-free library preparation using Hyper Library construction kit from Kapa Biosystems and subsequently sequenced on 300 cycle run (2×150nt reads) on two lanes of Illumina HiSeq 4000. This resulted in a total of 1.35 billion paired-end reads. Raw data for these 18 metagenomes was deposited in NCBI with bioproject number PRJNA552823. The reads were subject to adaptor removal and quality trimming using Trimmomatic (Bolger et al. 2014) which included adaptor removal, clipping of the first and last three bases, trimming of sequences where four-bp sliding window quality threshold was below Q20, and discarding of all trimmed sequences less than 75 bp in length. This resulted in a total of 1.03 billion paired end reads which were analyzed using a few different approaches (see below).

#### 2.3.1 Reference genomes based annotation

A total of 53 publicly available Genomes/genome assemblies for nitrifying organisms were downloaded from NCBI (RefSeq version 86) (Table S1). This included genome assemblies for 12 AOA, five anammox bacteria, 17 AOB, 10 comammox bacteria, and nine NOB. Reads from all samples were competitively mapped to the reference genomes using bwa (Li and Durbin 2010) followed by extraction of properly paired mapped reads (samtools view -f2) using SAMtools (Li et al. 2009). The extracted reads were then again competitively aligned against reference genomes using BLAST (Altschul et al. 1990) with a criterion of 90% sequence identity and 90% read coverage. The reads per kilobase million (RPKM) metric was used as a measure of the reference genome relative abundance in each sample and was estimated by dividing the number of reads aligning to reference genome at aforementioned criteria with the number of kilobases in each reference genome and the millions of reads per sample.

#### 2.3.2 SSU rRNA based community characterization

The paired end metagenomic reads were assembled using MATAM (Pericard et al. 2018) with the SILVA SSU rRNA (Release 132) database (Quast et al. 2013) as the reference for assembly of the 16S rRNA genes with a minimum 16S rRNA gene length threshold of 500 bp. The assembled 16S rRNA genes were classified against the RDP database (Cole et al. 2014) using Naïve Bayesian classifier approach (Wang et al. 2007). The relative abundance of each 16S rRNA gene was estimated by dividing the number of reads mapping to each MATAM assembled 16S rRNA gene with the total number of reads mapping to all assembled 16S rRNA genes per sample.

#### 2.3.3 Gene-centric de novo assembly

The paired end reads from each sample assembled into contigs using metaSpades (Nurk et al. 2017); where two sets of reads were available from attached and suspended phase from same bioreactor (i.e., JAMM, JAMS, DUR) or from two parallel operated bioreactors (GRE), these were co-assembled. MetaSpades assembly/co-assembly was carried out with kmers 21, 33, 55, 77, 99, and 127. The assembled contigs were filtered to remove all contigs smaller than 500 bp and subject to gene calling using prodigal (Hyatt et al. 2010) and annotation against the KEGG database (Kanehisa et al. 2016) using DIAMOND (Buchfink et al. 2015). Genes identified as encoding for *amoA* (KO:K10944), *amoB* (KO:K10945), and *nxrA* (KO:K00370) were extracted for further analyses. An analysis of genes assembled using *de novo* assembly approach for the complete metagenome revealed that several of the identified genes were highly fragmented. To circumvent this limitation, gene-centric *de novo* assembly was used. Specifically, paired end reads were mapped to a curated amino acid database of *amoA, amoB*, and nxrA genes using DIAMOND (Buchfink et al. 2015). The mapped reads were then assembled into contigs metaSpades (Nurk et al. 2017), followed by gene calling using prodigal (Hyatt et al. 2010) and annotation against the KEGG database (Kanehisa et al. 2016). The *amoA, amoB*, and *nxrA* genes identified using the complete and gene-targeted *de novo* assembly approach were combined, de-duplicated, and further curated by evaluating phylogenetic placement of assembled genes. Specifically, reference alignments for *amoA, amoB*, and *nxrA* genes were created by extracting corresponding genes from the reference assemblies and aligning with MUSCLE (Edgar 2004) followed by construction of a reference tree for each gene using RAxML (Stamatakis 2014). The *amoA, amoB*, and *nxrA* genes were placed on the gene-specific reference tree using pplacer (Matsen et al. 2010). Annotated genes that did not conform to known phylogeny of *amoA* and *amoB* for AOB, AOA, and comammox bacteria and *nxrA* genes for NOB and comammox bacteria were discarded. Finally, the relative abundance of the curated genes in each sample was estimated using the RPKM metric. Specifically, the sum of reads mapping to contig containing gene of interest per sample were divided by number of kilobases in for each contig and the millions of reads per sample.

#### 2.3.4 Recovery, phylogenomic placement, and annotation of Nitrospira metagenome assembled genomes

Assembled contigs from nine metagenomes corresponding to seven nitrogen removal systems where comammox bacteria were detected (i.e., DCWM, GRE1/GRE2, JAMM/JAMMSM, KIN, NEU, WIL, and YAN) were pooled. Reads from each metagenome were mapped to all contigs with bowtie2 (default parameters, version 2.1.0) (Langmead and Salzberg 2012) prior to binning with MetaBAT2 (-m 2000, version 2.12.1) (Kang et al. 2015). The completion and redundancy of the resulting bins were estimated with CheckM (lineage_wf, version 1.0.12) (Parks et al. 2015). Reads mapping to bins ≥ 50% complete were extracted and used to re-assemble the bin with Unicycler (default parameters, version 0.4.7) (Bankevich et al. 2012, Li et al. 2009, Wick et al. 2017). The quality and taxonomy of re-assembled bins were evaluated with CheckM (Parks et al. 2015) and the Genome Taxonomy Database Toolkit (GTDB-Tk 0.2.2, database release r86 v3) (Parks et al. 2018). Bins greater than 70% complete and classified as Nitrospira by GTDB-Tk were retained for manual refinement with Anvi’o (Eren et al. 2015). Following refinement, the quality and taxonomy of the bins were re-assessed with CheckM (Parks et al. 2015) and the Genome Taxonomy Database Toolkit (Parks et al. 2018), selecting metagenome assembled genomes (MAGs) with completeness and redundancy estimates of ≥ 70% and ≤ 10%, respectively. Open reading frames (ORFs) were predicted using Prodigal (Hyatt et al. 2010) v2.6.3 for all *Nitrospira* MAGs recovered from this study (n=10). KEGG orthologies (KO) were assigned to predicted ORF’s in these 10 genomes against 10,108 HMM models of prokaryota in KEGG(Kanehisa et al. 2016) database v90.1 using kofamscan (Aramaki et al. 2019). To investigate the phylogeny of these 10 *Nitrospira* MAGs recovered from this study, 32 previously publicly available N*itrospira* genomes were downloaded from Genbank (Table S2) (Poghosyan et al. 2019) and used as reference genomes for phylogenetic tree reconstruction. Phylogenomic tree reconstruction was conducted by Anvi’o (Eren et al. 2015) v5.5. ORFs were predicted for aforementioned 32 reference genomes and 10 MAGs from this study using Prodigal (version 2.6.3) (Hyatt et al. 2010) and then searched against a collection of HMM models summarized by Campbell et al. (Campbell et al. 2011) using hmmscan (version 3.2.1) (Eddy 2011) including 48 ribosomal proteins. Only genomes containing more than genes encoding for 40 of the 48 ribosomal proteins were included in downstream phylogenomic analyses. Alignments for each gene were conducted using muscle (Edgar 2004), alignments were concatenated, and finally phylogenomic tree was constructed using FastTree (version 2.1.7) (Price et al. 2010).

### 2.4 Quantitative PCR for total bacteria and nitrifying populations and design of primers targeting *amoB* gene of clade A comammox bacteria

PCR thermocycling and reaction mix conditions of previously published primer sets for qPCR-based quantification of 16S rRNA gene of total bacteria (Caporaso et al. 2011), AOB (Hermansson and Lindgren 2001) and *Nitrospira* (Graham et al. 2007) and ammonia monoxygenase subunit A (*amoA*) gene for AOB (Rotthauwe et al. 1997) are shown in Table 2. Further, qPCR assays targeting the *amoA* gene of comammox bacteria were conducted using previously published primer sets (Fowler et al. 2018, Pjevac et al. 2017) and additional comammox *amoA* primer sets designed a part of this study (Table 2). However, these assays either resulted in unspecific product formation and non-detection of comammox bacteria in samples where metagenomic analyses indicated presence of comammox bacteria. As a result, new primer sets targeting the *amoB* gene of clade A comammox bacteria (only clade A comammox bacteria were detected via metagenomic analyses in this study) were designed. To do this, the *amoB* genes assembled from the metagenomic data were combined with *amoB* gene sequences from previous studies (Daims et al. 2015, Palomo et al. 2016, Pinto et al. 2015, van Kessel et al. 2015) and two primer sets were designed (Table 2). These new set of primers were tested with samples that were positive and negative for comammox bacteria, along with DNA extracts from *Ca* Nitrospira inopinata as positive control involving variation in annealing temperature, primer concentration, and template concentration.

**Table 2:**
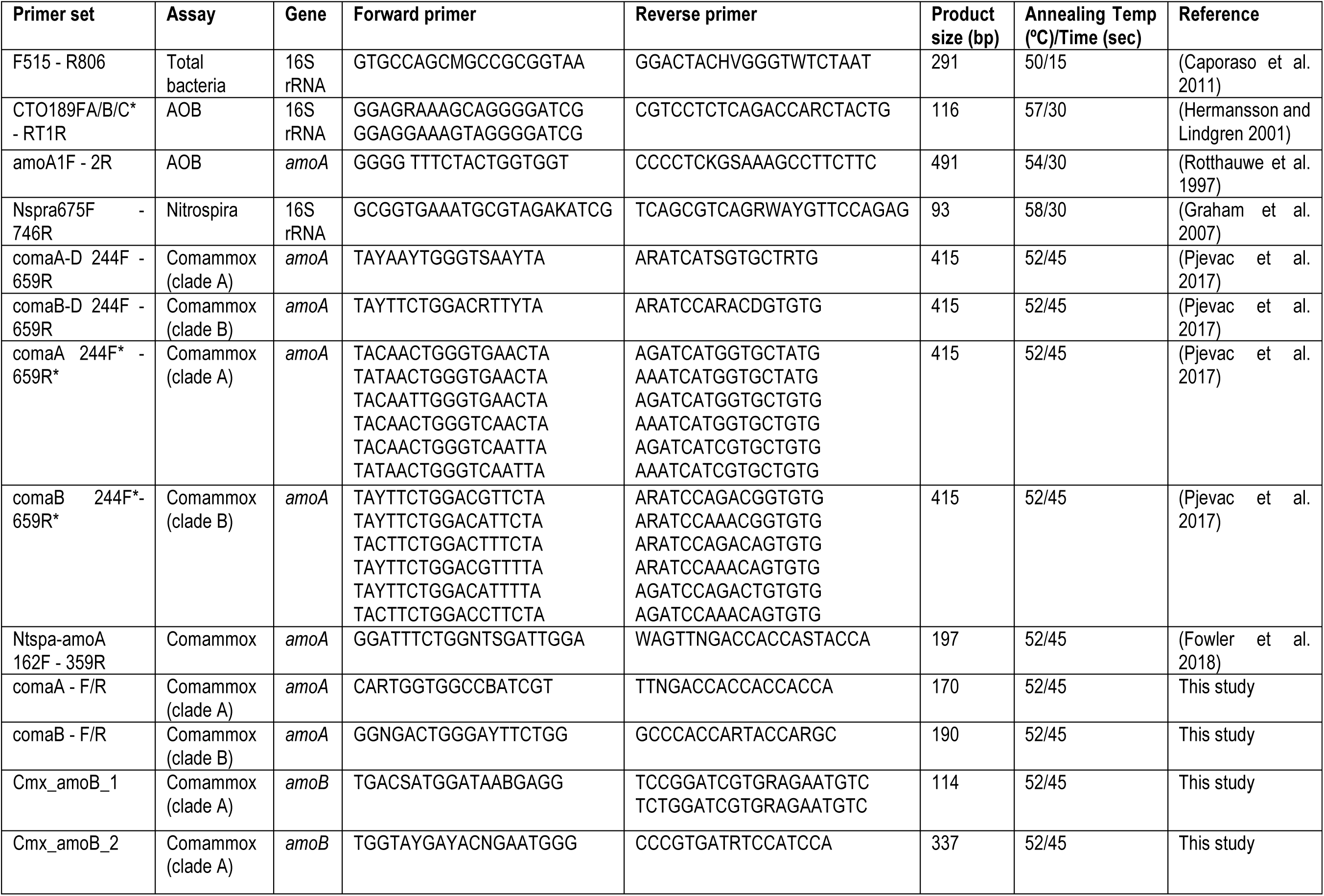
A summary of qPCR primers used and tested as part of this study (*equimolar proportions of forward and/or reverse primers if multiple primers are used).

The qPCR assays were performed on a QuantStudio 3 Real-Time PCR System (ThermoFisher Scientific, Cat. No. A28567) in 20 µL reaction volume including: 10 µL Luna Universal qPCR Master Mix (New England Biolabs, Inc., Cat. No. NC1276266), primers listed in Table 2, 5 µL of 10 times diluted DNA template and the required volume of DNAse/RNAse-Free water (Fisher Scientific, Cat. No. 10977015) to reach 20 µL reaction. Reactions were prepared by the epMotion M5073 liquid handling system (Eppendorf, Cat. No. 5073000205D) in triplicate. The cycling conditions were as follows: initial denaturing at 95°C for 1 min followed by 40 cycles of denaturing at 95°C for 15 s, annealing temperatures and times listed in Table 2, and extension at 72°C for 1 min. Melting curve analyses was performed at 95°C for 15 s, 60°C for 1 min, and 95°C for 15 s. A negative control (NTC) and a standard curve ranging from 10^3^-10^9^ copies of 16S rRNA gene of *Nitrosomonas europaea* for total bacteria assay and 10^2^ – 10^8^ copies of 16S rRNA genes for *Nitrosomonas europaea* and *Ca* Nitrospira inopinata for the AOB and *Nitrospira* assays, respectively and 10^2^ – 10^8^ copies for *amoA* of *Nitrosomonas europaea* and *Ca* Nitrospira inopinata for the AOB and comammox assays, respectively and *amoB* gene of *Ca* Nitrospira inopinata for the comammox assays were included in the qPCR analysis.

### 2.5 Statistical analyses

Statistically significant differences in the abundance of nitrifying organisms between nitrogen removal systems was evaluated using the non-parametric Kruskal-Wallis or Wilcoxon rank sum test in R as appropriate (RCoreTeam 2014). Correlations between any two variables were determined using Spearman rank correlations and BIOENV analysis within the R package “vegan” to identify process and environmental variables that demonstrate significant correlation with changes in nitrifier population abundances. All statistical tests and figure generation (Wickham 2009) were performed in R (RCoreTeam 2014).

## 3.0 Results and Discussion

### 3.1 Overview of nitrogen removal systems included in this study

As part of this study, we sampled fourteen full-scale nitrogen removal systems of varying process configurations. Specifically, we sampled nine nitrification-denitrification (ND), two simultaneous nitrification-denitrification (SND) mainstream systems, and three partial nitrification-anammox (PNA) sidestream systems with diverse process subtypes (Table 1). The process sub-types ranged from multi-stage suspended growth systems with secondary clarification to sequencing batch reactors (SBRs) ones to membrane bioreactors (KIN) to systems with significant attached growth components (e.g., DUR, JAMM, JAMS). The typical process parameters for each of these systems is shown in Figure 1. All nitrogen removal systems included in this study performed normally during this study with no significant and sustained process upset.

**Figure 1:**
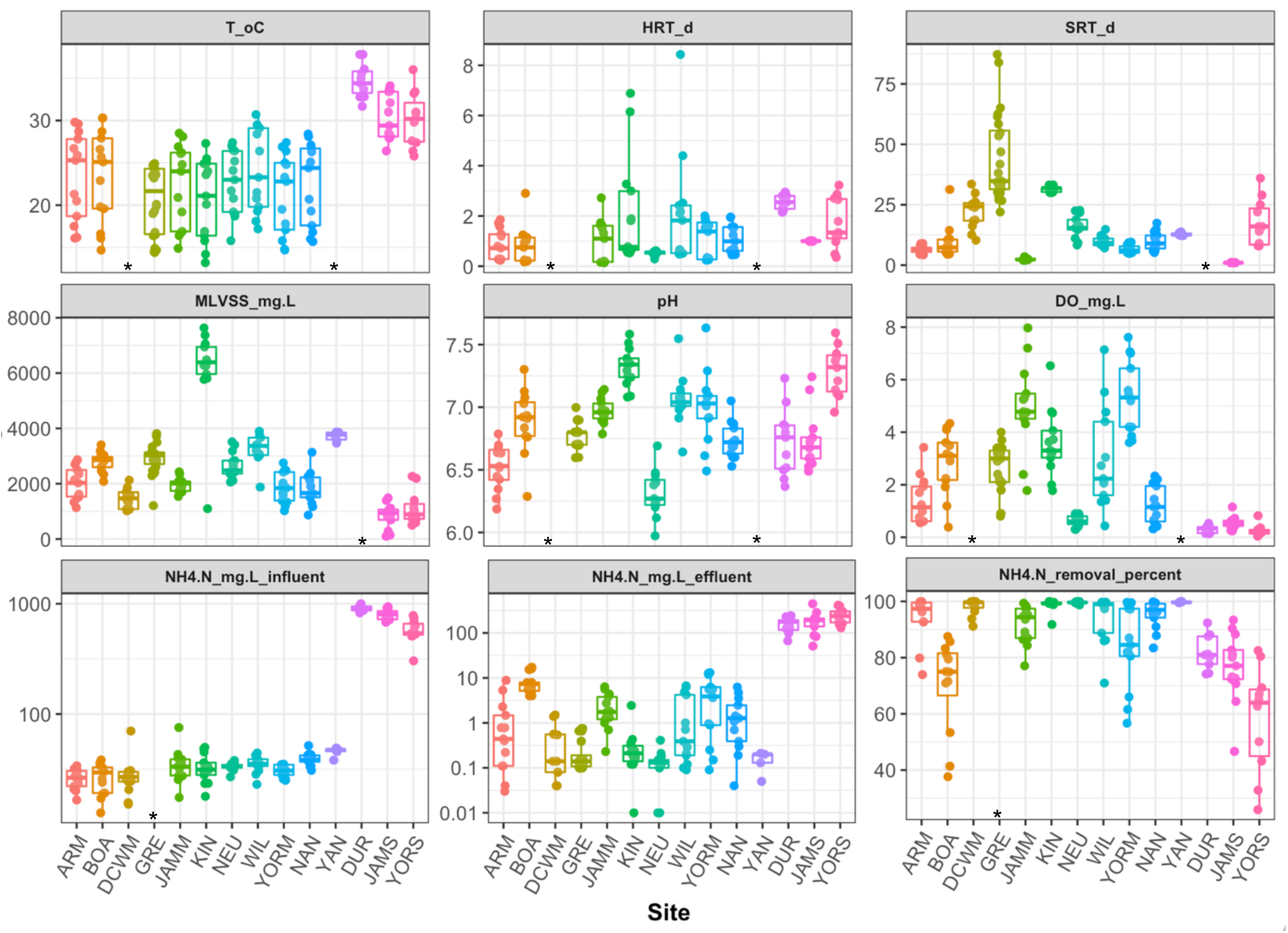
Overview of key process parameters and performance metrics monitored as part of this study for 14 nitrogen removal systems. Column indicated with “*” indicates lack of relevant process data for the respective nitrogen removal system.

### 3.2 Identification of nitrifying populations using reference genome mapping and 16S rRNA gene assembly

Mapping of reads to reference genomes indicated that *Nitrospira*-like bacteria were dominant members of the nitrifying communities in all ND and SND systems followed by *Nitrosomonas*-like bacteria, with no or very low detection of AOA (Figure 2A). Nearly all detected AOB belonged to the genus *Nitrosomonas* with low levels of detection of AOB within the genera *Nitrosococcus* and *Nitrosospira* in two systems (i.e., ARM and BOA). Metagenomic reads mapped to comammox references genomes within the genus *Nitrospira* for samples from six ND and one SND system, with abundances higher than canonical *Nitrospira*-NOB for three ND systems (i.e., GRE, KIN, and NEU) and one SND system (i.e., YAN). Genera containing other canonical NOB, i.e. *Nitrotoga* and *Nitrobacter*, were detected in two ND and one SND system with *Nitrotoga* being the primary NOB in BOA. Mapping of metagenomic reads from PNA systems to reference genomes primarily resulted in the detection of anammox bacteria (genus: *Brocadia*) and AOB (genus: *Nitrosomonas*), with no detectable presence of *Nitrospira*. It is important to note that while metagenomic reference genome based approach may be useful for detection of organisms, its quantitative value is limited. This limitation likely emerges from the reliance of a reference database of genomes and the stringent criteria used for read mapping (i.e., 90% sequence identity between metagenomic read over 90% of the read length). Thus, it is possible that the significant over representation of *Nitrospira* in the metagenomic reference genome-based approach emerges from a high level of genomic similarity between *Nitrospira* in the samples and in the reference database and under representation of AOB emerges from a low level of genomic similarity between AOB in the samples and in the reference database. This is the most likely scenario because genomic similarity between organisms within the same genus can be as low as 75% (Jain et al. 2018).

**Figure 2:**
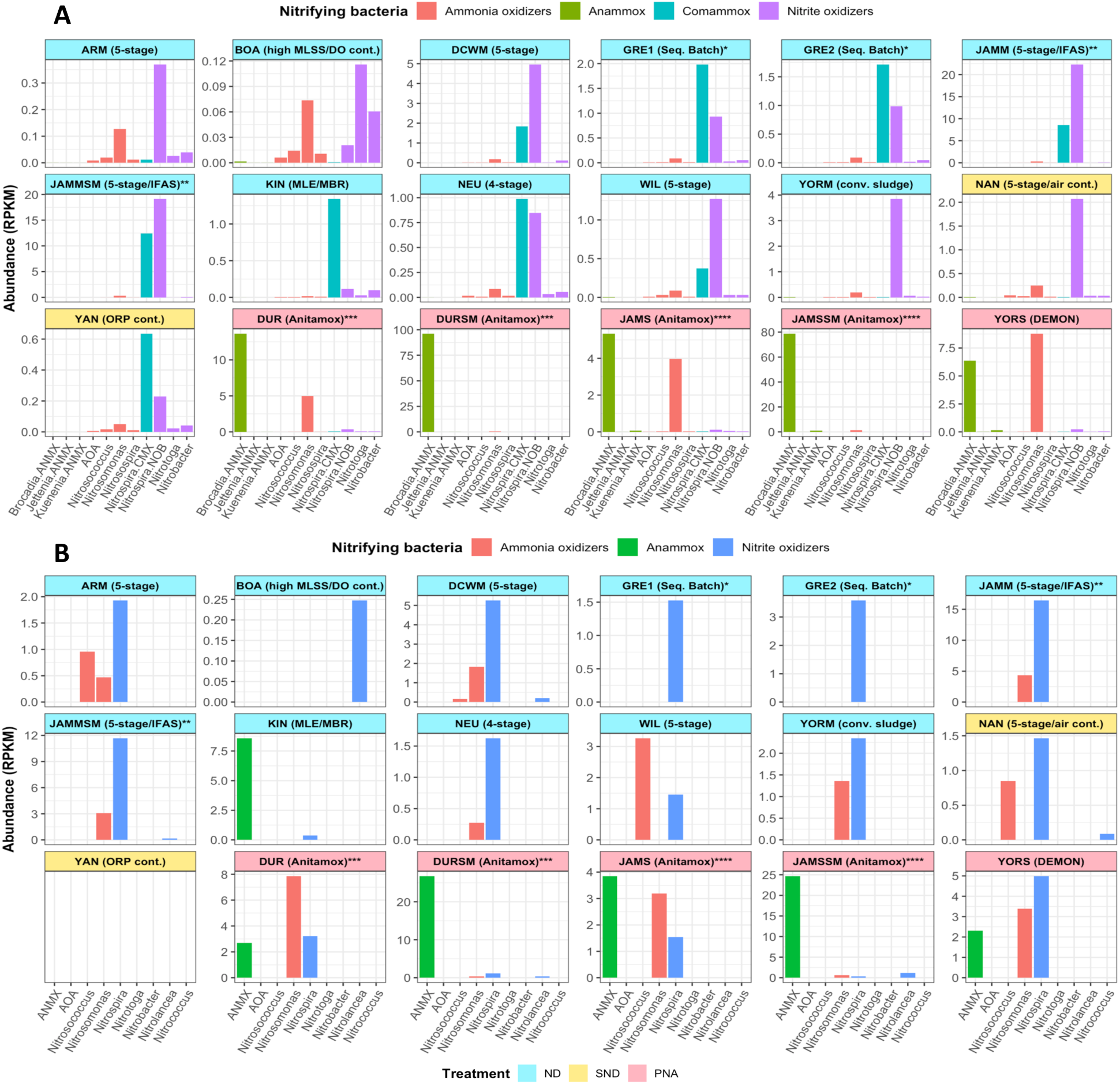
Relative abundance of nitrifying organisms in nitrogen removal systems based on (A) mapping of reads to reference genomes and (B) classification of MATAM assembled 16S rRNA genes. Bar colors indicate type of nitrifying organism detected (see color legend below each figure), while facet color indicates process type (i.e., red: PNA, Blue: ND, and Yellow: SND). GRE1/GRE2* panels present abundance of nitrifying populations from two different reactors at the same treatment plant. JAMM/JAMMSM**, DUR/DURSM***, and JAMS/JAMSSM**** panels present abundance of nitrifying populations in the suspended phase and attached growth phase of the system, respectively.

In contrast, a subset of genes (particularly ribosomal genes) may be far more reliable for relative quantitation of microbial abundance within the larger community. Thus, we used MATAM to identify reads originating from the 16S rRNA gene within each metagenomic sample and subsequently assemble them and finally determine the relative abundance of each gene in the sample microbial community. The reference genome mapping results were largely consistent with classification of MATAM-assembled 16S rRNA genes (Figure 2B) with *Nitrospira*- and *Nitrosomonas*-like bacteria dominant in ND and SND systems and *Brocadia*- and *Nitrosomonas*-like bacteria being dominant in PNA systems. Three of the 14 systems included in this study consisted of distinct suspended and attached growth phase, i.e., ND IFAS system (i.e., JAMM) and two side-stream PNA systems (DUR and JAMS). Relative abundances of *Nitrospira*- and *Nitrosomonas*-like bacteria were similar between the suspended and attached growth phase for JAMM while *Brocadia*-like bacteria were enriched in the attached growth phase for DUR and JAMS. The key differences between reference genome- and 16S rRNA gene analyses were for the PNA systems and one ND system (i.e., BOA). Specifically, *Nitrospira*-like bacteria were also detected in all PNA system included in this study using 16S rRNA gene analyses compared to reference mapping of metagenomics reads and while *Nitrolancea*-like bacteria were the primary nitrifying bacteria in BOA, 16S rRNA gene analyses suggested high abundance of *Brocadia*-like bacteria in KIN as well. The discrepancies between reference genome based analyses and 16S rRNA gene assembly based analyses may result from the limitation of the prior (as stated above), inefficient assembly of 16S rRNA genes due to highly conserved regions (Miller et al. 2011, Pericard et al. 2018), or a combination of both.

### 3.3 Functional gene based metagenomic identification of ammonia and nitrite oxidizing bacteria

Mapping of reads to curated *amoA, amoB*, and *nxrA* genes, followed by gene-centric *de novo* assembly was used to detect and estimate the relative abundance of AOB, comammox bacteria, and NOB in the metagenomic datasets from the 14 systems. Subsequently, all assembled and annotated genes were placed on gene specific phylogenetic trees to eliminate potentially misannotated *amoA, amoB*, and *nxrA* genes. This analysis revealed that nearly all annotated AOB belonged to *Betaproteobacteriales* (Figure 3A) and contained representative sequences from ND, SND, and PNA systems, while *amoA* and *amoB* sequences from comammox bacteria were detected in DCWM, GRE, JAMM/JAMMSM, KIN, NEU, WIL, and YAN (Figure 3B). These results are qualitatively consistent (i.e., presence/absence of comammox bacteria) with those obtained by reference genome based analyses of comammox bacteria. The relative abundance of comammox bacteria was higher than AOB in JAMM/JAMMSM, while both were equally abundant two ND systems (i.e., GRE and NEU) and one SND system (i.e., YAN). In contrast, comammox were less abundant than AOB in DCWM and WIL; both of these are ND systems. Interestingly, no AOB were detected in KIN (an MBR/MLE system) with comammox being the only ammonia oxidizer. The results for KIN were consistent with that of reference genome based and 16S rRNA gene based analyses, where no AOB were detected. Further, the similarity in relative abundance of *amoA, amoB*, and *nxrA* genes indicates that all detected *Nitrospira* bacteria in this system are likely to be comammox bacteria. The non-detection of other AOB (e.g., *Nitrosospira, Nitrosococcus*) and NOB (e.g., *Nitrotoga*) that were detected by reference genome based or 16S rRNA gene assembly based analyses could be due to challenges with *de novo* assembly genes from low abundance microorganisms (i.e., insufficient sequencing depth and thus low coverage).

**Figure 3:**
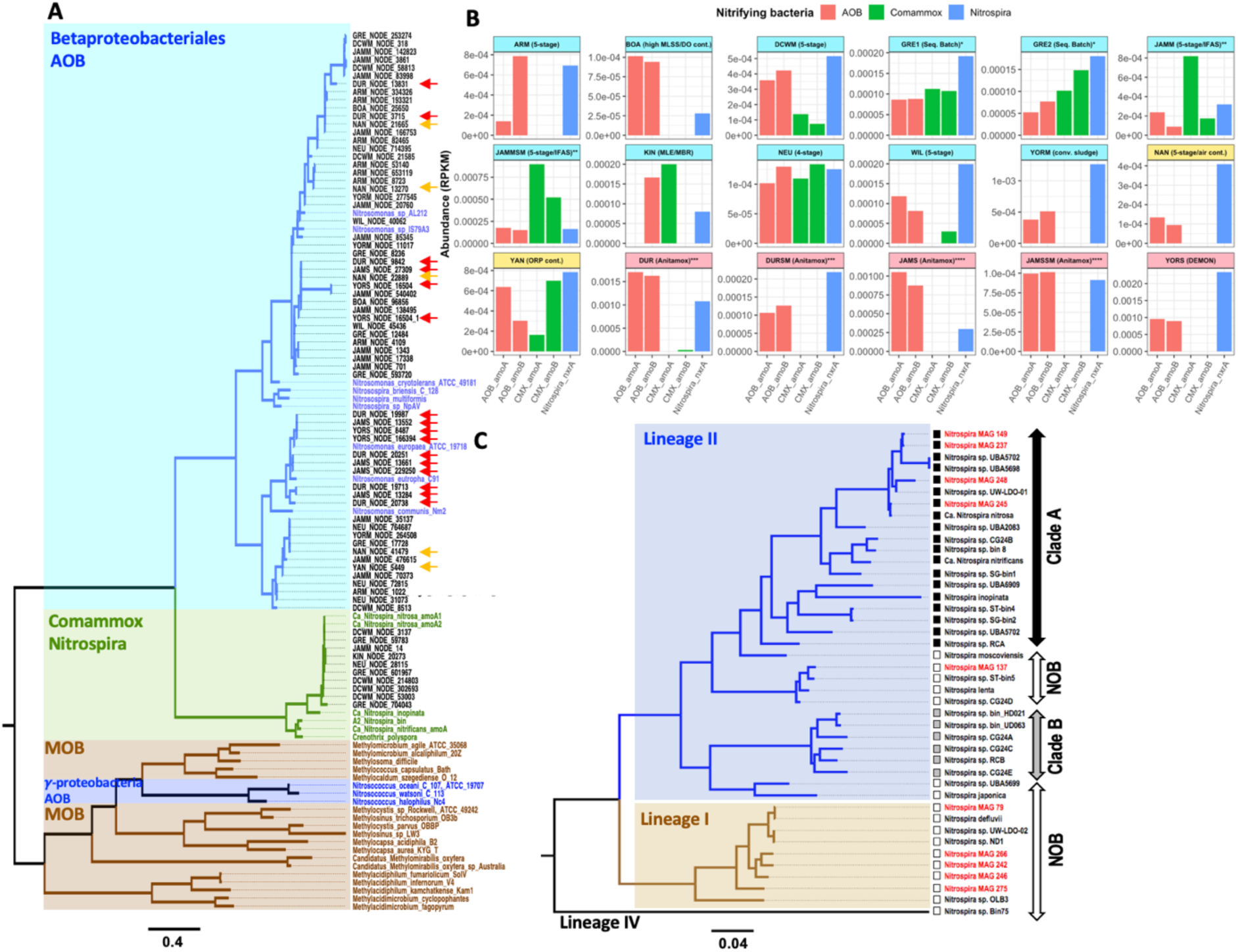
(A) Maximum likelihood phylogenetic tree of *amoA* genes recovered from gene centric *de novo* assembly along with reference *amoA* and *pmoA* gene sequences. Clades are colored by functional grouping and labels for reference sequences within each cluster are colored according to the functional group, while amoA sequences recovered from this study are shown in black labels. Red and orange arrows indicate amoA gene sequences recovered from PNA and SND systems respectively, while those recovered from ND systems are not annotated. amoA gene sequences from YAN and KIN are not shown in the phylogenetic tree due to recovery of short sequences not suitable for tree construction. (B) Abundance (RPKM) of amoA, amoB gene sequences of AOB and comammox bacteria, and nxrA gene sequences of Nitrospira are shown. (C) Phylogenetic placement of Nitrospira MAGs (red label) with 32 reference genomes (black label). Clade A (black squares), clade B (grey squares) comammox bacteria and canonical NOB (open squares).

A consistent aspect of systems with prevalent comammox populations was their high total solids retention time (SRT). For instance, KIN is an MBR system with an estimated SRT of ∼30 days, DCWM is a ND system with a total SRT of ∼30 days, and GRE is an SBR system with an SRT > 30 day. While JAMM/JAMMSM (i.e., James River) has an SRT of 2.5 days for the suspended phase, this is an IFAS system with a significant attached growth component. This suggests that irrespective of the process configuration type (ND, SND) and/or mode of operation (continuous, SBR, attached growth), the key similarity among comammox prevalent systems is their high SRT or presence of attached growth component (i.e., implicitly high SRT). This is consistent with thermodynamic and metabolic pathway modeling that suggests that comammox bacteria have a very low specific growth rates compared to AOB and NOB (Costa et al. 2006). For instance, recent literature suggests that comammox maximum specific growth rates are 3-10, 5-11, and 2 times lower than that of Nitrospira-NOB, AOB, and AOA (Kits et al. 2017, Lawson and Lücker 2018). While this explains the prevalence of comammox in long SRT system, it does not explain the absence of AOA in these systems. While AOA are highly prevalent in other ecosystems (i.e., soils, drinking water, marine environments) (Prosser and Nicol 2008), they are rarely detected in wastewater systems (Chao et al. 2016, Pjevac et al. 2017) including those in this project. It is plausible that the absence of AOA and presence of comammox may be explained by the difference in their affinity for ammonia, i.e., AOA with K_m_ (NH_3_) ranging from 0.36 to 4.4 μM compared with reported *Ca* Nitrospira inopinata K_m_ (NH_3_) of 0.049 μM (Kits et al. 2017, Lawson and Lücker 2018).

All comammox based *amoA* and *amoB* gene sequences detected using metagenomic sequencing closely clustered with *Ca* Nitrospira nitrosa and distinct from other clade A bacteria (e.g., *Ca* Nitrospira nitrificans, *Ca* Nitrospira inopinata) (Figure 3B). Further, all Nitrospira MAG’s recovered from the genome binning process were associated with either lineage 1 or 2 *Nitrospira* and included six canonical NOB and four clade A comammox. Genome statistics for the recovered MAGs are shown in Table 3. Consistent with previously described clade A comammox bacteria, all comammox MAGs contained some or all genes associated with both ammonia and nitrite oxidation, except for *Nitrospira* MAG 248 which is likely due to lower completeness (i.e., 70%). Similarly, these comammox MAG’s contained genes involved in urea uptake and hydrolysis - a feature shared with the other recovered canonical *Nitrospira* NOB MAGs. Further, similar to other clade A comammox bacteria, the recovered comammox MAG’s lacked ammonium transporter (Amt) or any of formate dehydrogenase genes which are reported present in clade B comammox as well as canonical NOB within lineage II *Nitrospira* (Poghosyan et al. 2019). Finally, similar to previously reported clade A comammox bacteria, the comammox MAG’s recovered in this study also indicated presence of genes associated with group 3b [NiFe] hydrogenase.

**Table 3:** Statistics of the Nitrospira MAG’s assembled in this study and their functional grouping.

| ***Nitrospira* MAGs** | **79** | **137** | **149** | **237** | **242** | **245** | **246** | **248** | **266** | **275** |
| --- | --- | --- | --- | --- | --- | --- | --- | --- | --- | --- |
| *Nitrospira* lineage | 1 | 2 | 2 | 2 | 1 | 2 | 1 | 2 | 1 | 1 |
| Completeness (%) | 94.94 | 87.92 | 75.23 | 93.3 | 89.09 | 82.17 | 84.94 | 70.99 | 83.58 | 97.67 |
| Redundancy (%) | 1.36 | 9.15 | 1.14 | 3.86 | 2.36 | 4.6 | 4.09 | 2.12 | 2.73 | 7.78 |
| Genome size (Mbp) | 3.49 | 4.07 | 3.08 | 4.54 | 4.02 | 3.8 | 3.78 | 2.59 | 3.73 | 3.91 |
| GC content | 59.14 | 58.05 | 54.87 | 54.83 | 58.41 | 54.64 | 58.36 | 54.86 | 58.42 | 58.79 |
| Number of contigs | 136 | 410 | 229 | 48 | 114 | 203 | 98 | 412 | 78 | 54 |
| N50 of contigs | 40833 | 25623 | 21078 | 171941 | 177531 | 34975 | 85503 | 8266 | 78457 | 794701 |
| genes | 3361 | 4115 | 3141 | 4498 | 3886 | 3779 | 3694 | 2693 | 3625 | 3753 |
| Protein coding genes | 3322 | 4062 | 3113 | 4455 | 3845 | 3743 | 3653 | 2668 | 3583 | 3708 |
| rRNA | 0 | 1 | 1 | 3 | 0 | 0 | 0 | 0 | 0 | 0 |
| tRNA | 38 | 51 | 26 | 40 | 41 | 36 | 41 | 25 | 41 | 44 |
| Functional group | NOB | NOB | CMX | CMX | NOB | CMX | NOB | CMX | NOB | NOB |

### 3.4 Development of qPCR assay for quantitative detection of comammox bacteria

We used previously developed primer sets (Fowler et al. 2018, Pjevac et al. 2017) to quantify the abundance of comammox bacteria in samples collected as part of the study (Table 2). Pjevac et al. (2017) provided two different primer sets targeting the *amoA* gene both clade A and clade B comammox bacteria; one primer set was the degenerate primer set while the other contained an equimolar proportion of six forward and six reverse primers. In contrast, Fowler et al. (2017) developed a single primer set targeting the *amoA* gene of both clade A and clade B comammox bacteria. While both primer sets showed excellent PCR efficiency with DNA extracts from pure culture of *Ca* Nitrospira inopinata, efforts to PCR amplify the *amoA* gene from samples included in this study demonstrated either non-detection of comammox bacteria in the six systems with metagenomic evidence of their presence or unspecific product formation (Figure 4A, B). This was also consistent for *amoA* specific primers designed in this study (Table 2). The same issue was also highlighted by Beach and Noguera, (2019), who suggested the use of species primers for comammox detection based on the system of interest (i.e., wastewater, drinking water, etc). The comammox *amoA* primers were also tested against the comammox *amoA* genes recovered from the metagenomic sequencing data. This *in silico* analysis indicated that primer sets comaA-D (244f-659R), comaA (244F-659R), and comA-F/R matched recovered *amoA* genes, yet the issue with unspecific product amplification and non-detects in some samples were persistent despite additional efforts to optimize annealing temperatures, template concentration and PCR additives (i.e., DMSO, BSA, magnesium, etc). Therefore, we developed new primer sets targeting the *amoB* gene of comammox bacteria, specifically clade A comammox bacteria; the primary comammox clade detected in our study and in wastewater treatment systems by recent studies (Chao et al. 2016, Pjevac et al. 2017, Roots et al. 2019, Spasov et al. 2019). The *amoB* gene exhibits phylogenetic clustering consistent with that of *amoA* gene and places comammox bacteria in a distinct cluster from all other ammonia oxidizers. Thus, we developed two primers sets targeting the *amoB* gene of comammox bacteria with an expected product size of 114 and 337 bp. Both primer sets resulted in high specificity (i.e., no unspecific amplification) (Figure 4C), high PCR efficiency (∼94%) (Figure 4D), and a clean melting curve (Figure 4E). Further, the qPCR bases estimates of the ratio of comammox bacteria (determined using *amoB* specific primers) to total ammonia oxidizers was highly correlated for the ratio of comammox bacteria to total ammonia oxidizers determined using metagenomic data involving gene centric *de novo* assembly (Figure 4F) and mapping of metagenomic reads to reference genomes (Figure 4G). This suggests that the newly designed primers are specific to clade A comammox bacteria and accurately capture their abundance in complex nitrifying communities.

**Figure 4:**
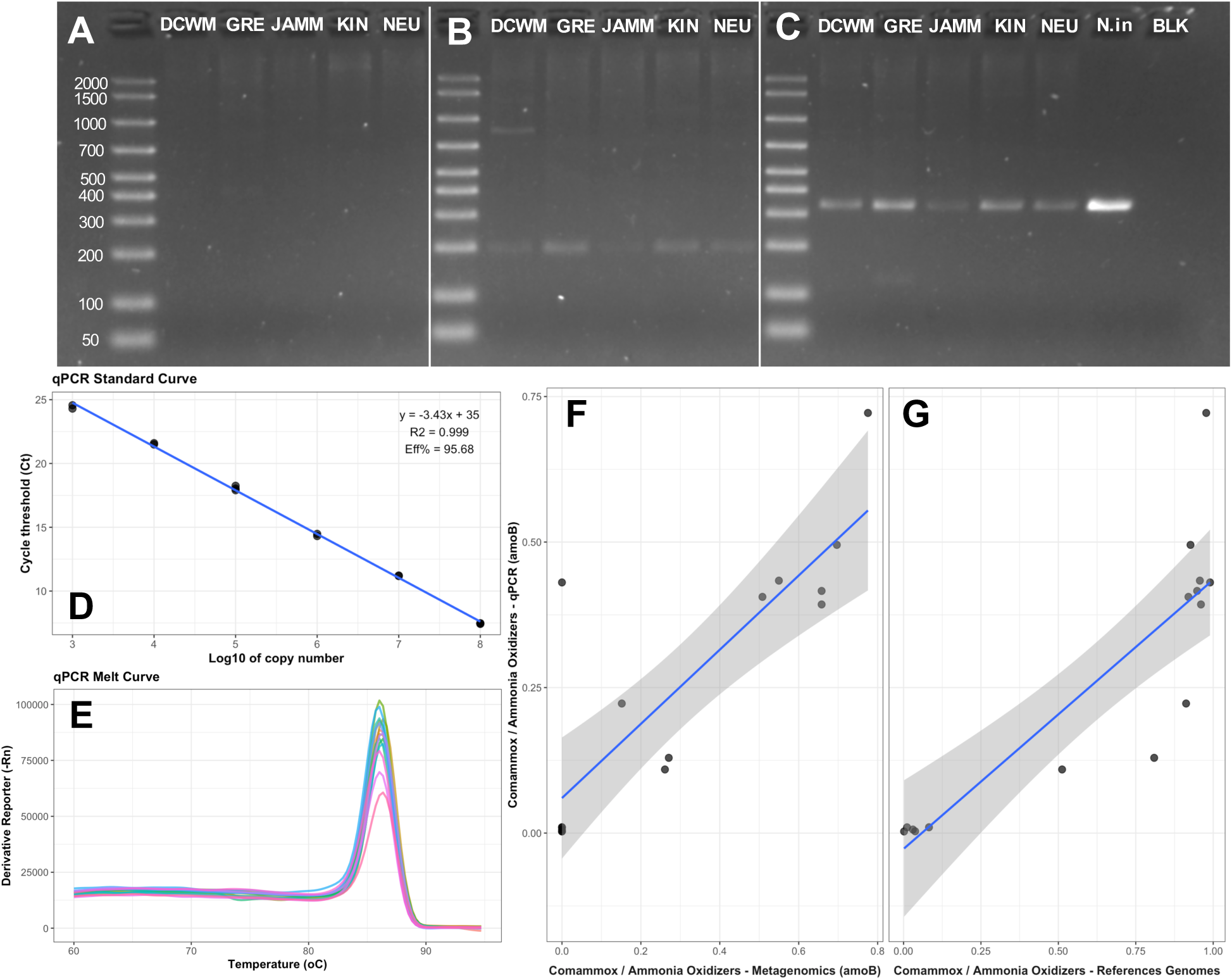
Previously published primer sets for the amoA gene of comammox bacteria, i.e. (A) comaA, and (B) Ntspa-amoA, result in non-detection and unspecific amplification samples from DCWM, GRE, JAMM, KIN, NEU where comammox bacteria were detected using metagenomics. In contrast the newly designed *amoB* specific primers (C) result in a single band formation of the right size for these samples which is consistent with that amplified from Ca Nitrospira inopinata (N.in). BLK stands for PCR negative control. The primers exhibit excellent PCR efficiency (D), demonstrate clean melting curves (E), and provide estimates of comammox bacterial abundance consistent with those seen in PCR-free metagenomics analyses (F, G).

### 3.5 Quantitative detection of nitrifying populations including comammox bacteria

Initial assessment of qPCR results from the *amoA* assay for AOB underestimated AOB abundance when compared to the qPCR data targeting the 16S rRNA gene for AOB (Supplemental Figure S1). In contrast, 16S rRNA gene based qPCR estimates for AOB were consistent with those obtained using metagenomics (Supplemental Figure S1). This issue was also reported by Dechesne et al. (2016) (Dechesne et al. 2016) who indicated that a*moA* primer sets do not provide sufficient coverage for AOB, while the 16S rRNA primer sets likely capture some non-AOB sequences as well. Our analyses suggest that while *amoA* primer sets did not provide sufficient coverage for AOB in our study, comparisons with metagenomic data indicated no evidence on unspecific non-AOB detection with 16S rRNA gene primers. As a result, all subsequent measurements of AOB were conducted using 16S rRNA gene based assays.

Canonical AOB and *Nitrospira* (including comammox and NOB) were detected in all systems irrespective of the nitrogen removal process configuration. The relative abundance of both groups ranges from 0.25 to 9% of total bacteria with an average of 1.71 and 1.65%, respectively, except for *Nitrospira* in the attached growth of JAMM/JAMMSM which reached a maximum relative abundance of 20% with an average of 8.56% (Figure 5A, B). For several systems, the abundance of *Nitrospira* in proportion to AOB was significantly and consistently above what would be expected if nitrification was being driven by AOB and NOB alone. These included the following ND systems: DCWM, attached growth phase of JAMM (i.e., JAMMSM), KIN, SND system: YAN, and PNA systems: attached growth phase of JAMS (i.e., JAMS). While comammox bacteria were detected in most systems with higher proportional abundance of *Nitrospira* over AOB (except for the PNA system: JAMS), the presence of comammox bacteria was not exclusive to them (Figure 5B). For instance, comammox bacteria were also detected in GRE and NEU where the abundance of *Nitrospira* and AOB were largely in line with expected proportions if nitrification was primarily driven by AOB and NOB (Costa et al. 2006). Comammox bacteria were not detected in four ND systems (ARM, BOA, NAN, YORM), one SND system (NAN), and the three PNA systems (DUR/DURSM, JAMS/JAMSSM, YORS), which was consistent with metagenomic observations. In all, comammox bacteria were detected in six ND systems (i.e., DCWM, GRE, NEU, WIL, KIN, JAM/JAMSM) and one SND system (i.e., YAN). The abundance of comammox bacteria in these systems ranged from 0.5% to ∼3% of total bacteria in the suspended growth phase, while it was as high (as high as ∼17%) in the attached phase of the IFAS systems (i.e., JAMMSM). In systems that included only suspended growth phase, comammox bacteria constituted between 20-70% of all ammonia oxidizers (i.e., AOB + comammox) for the duration of the study, with abundances equal to that of AOB in two ND systems (i.e., GRE and NEU). The proportion of comammox bacteria to that of total ammonia oxidizers varied significantly for two systems with low dissolved oxygen, i.e., KIN (ND) and YAN (SND), from 1 to 70% and at times significantly surpassed that of AOB.

**Figure 5:**
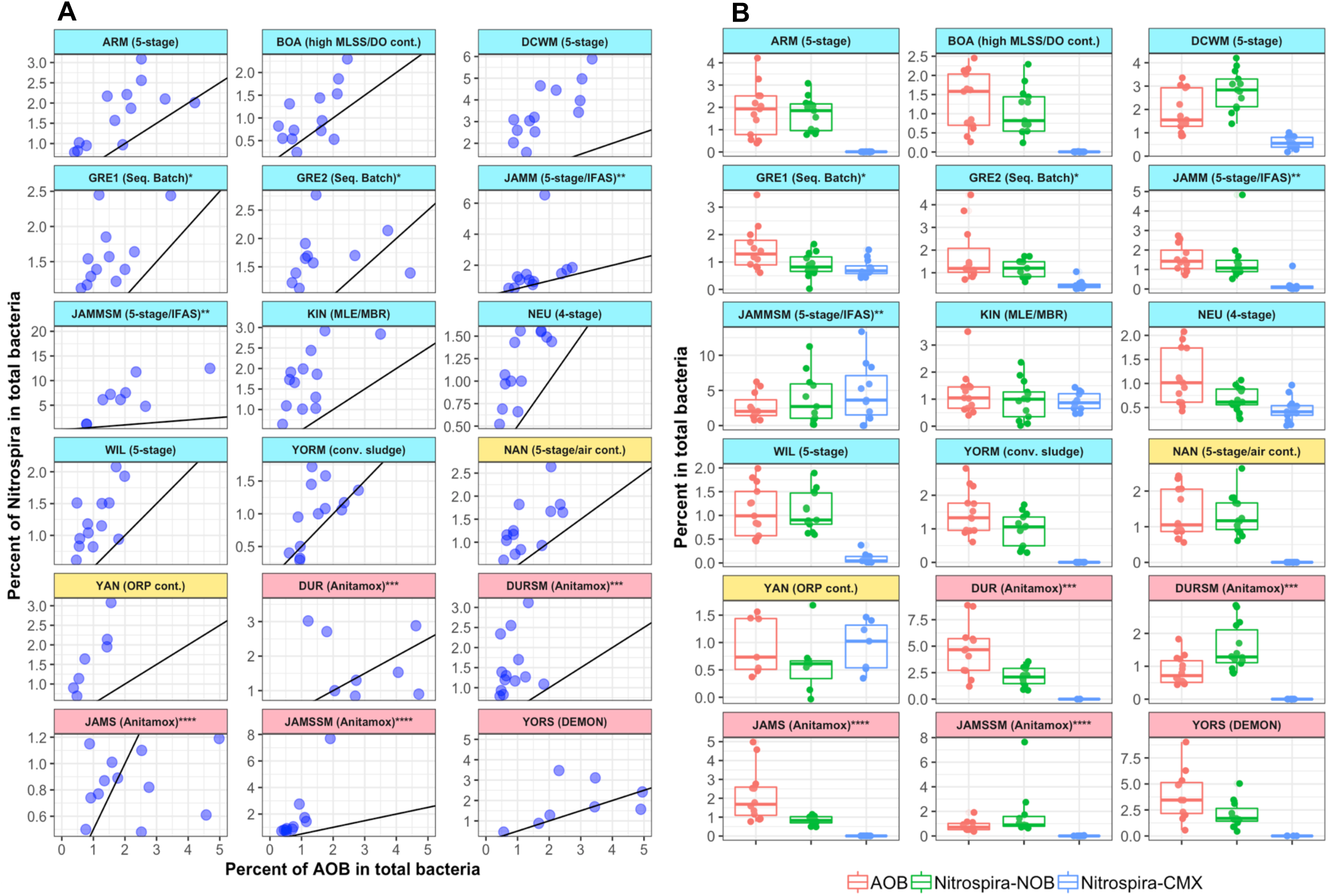
(A) Comparison of the abundance of Nitrospira (% of total bacteria) to that of AOB (% of total bacteria) indicates that several systems with higher proportional abundance of Nitrospira when compared to the theoretical estimates (black line) if nitrification was being driven by AOB and NOB. (B) The abundance of AOB, Nitrospira-NOB, and comammox bacteria as a proportional of total bacterial abundance indicates that comammox bacteria were (1) consistently detected in systems with metagenomic evidence of their presence, (2) their abundance was lower than AOB in suspended growth systems where they were detected and high in systems with attached growth phase. Asterisks levels indicate samples from the same system.

### 3.6 Comammox bacterial abundance was strongly associated with solids retention time

Comammox bacteria were prevalent and temporally persistent in systems with long SRTs, despite fluctuations in SRT levels within each system. The abundance of comammox bacteria as a proportion of all ammonia oxidizers (i.e., comammox + AOB) increased with increasing median SRT of the system (Figure 6A, 6B), with high abundances on the attached growth phase of the IFAS systems at JAMM. Further, the mean ammonia removal was typically higher in systems with comammox bacterial present compared to those without comammox bacteria (Wilcoxon rank sum test, p<0.0001) (Figure 6C). And finally, while comammox bacterial abundance increased with SRT (Figure 6F) (Spearman’s R = 0.52, p <0.0001), the abundance of *Nitrospira*-NOB and AOB did not exhibit significant association with SRT (Figure 6D, E). This suggests that increases in comammox abundance may not be associated with a concomitant decrease in AOB or NOB concentrations.

**Figure 6:**
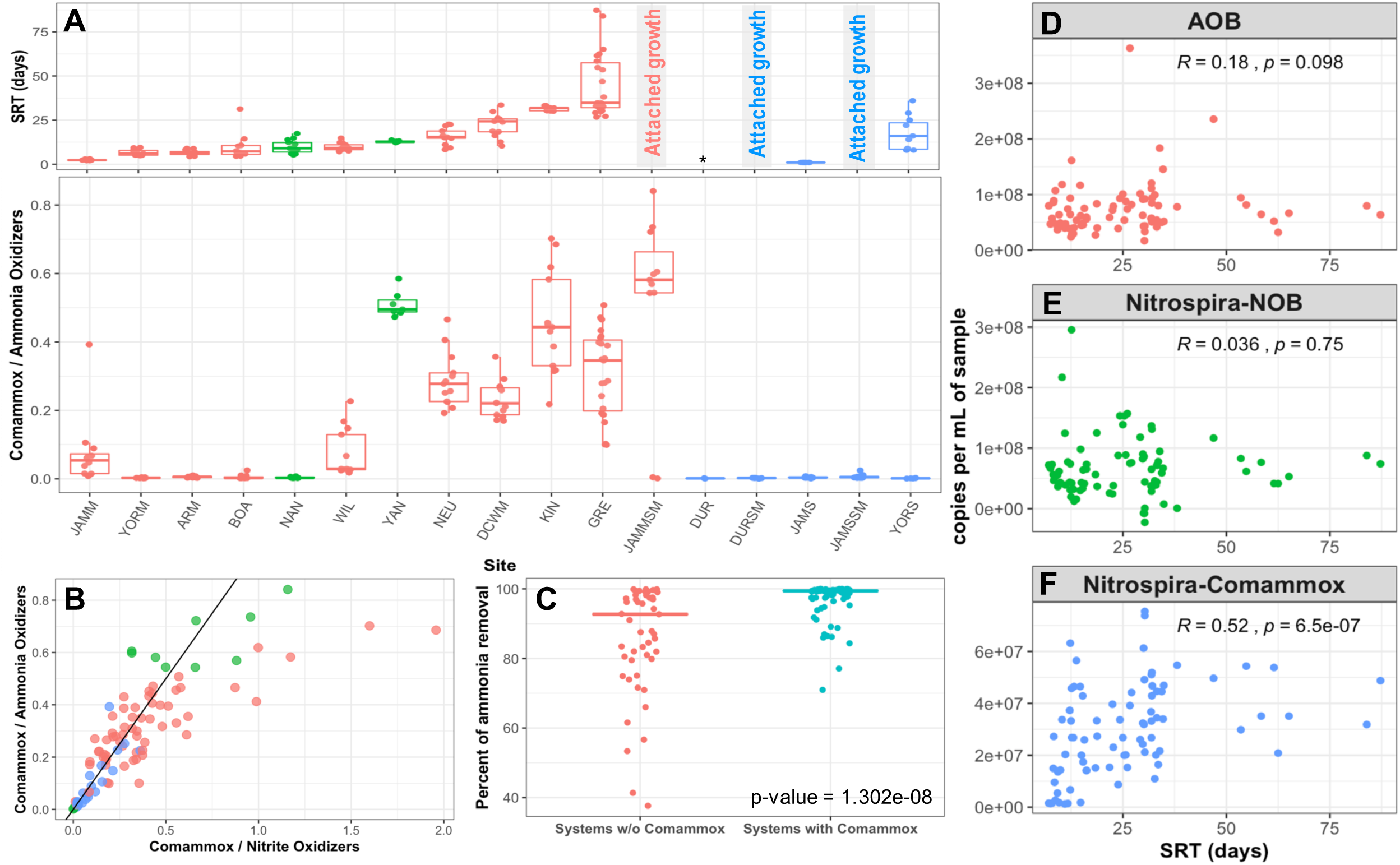
Increase in solids retention time (A) was associated with comammox bacteria constituting a greater proportion of all ammonia oxidizers (B). (C) Systems with comammox bacteria exhibited greater ammonia removal efficiencies compared to systems without comammox bacteria. This comparison does not include PNA systems. In contrast to AOB (D) and NOB (E), whose abundance was not associated with SRT, comammox bacterial abundance (F) was significantly and positively correlated with system SRT.

These data provide two potential insights into comammox bacterial relevance in nitrogen removal systems and their potential competitive dynamics or lack thereof with canonical nitrifiers. First, as stated earlier (section 3.3), comammox bacteria are preferentially enriched in systems with long SRT’s and systems with an attached phase component. While the prevalence in long SRT systems may be explained by their slower net growth rates compared to canonical AOB and NOB (Kits et al. 2017, Lawson and Lücker 2018), this does not provide basis for explaining their preferential enrichment over AOB and NOB with increasing SRT. It is also important to note that the AOB and NOB abundances were not associated with a concomitant increase in comammox bacterial abundance suggests that these bacteria may occupy exclusive niches within the nitrifying consortium. For instance, AOB and comammox may occupy independent niches at different ammonia concentrations due to different affinity levels for ammonia (Kits et al. 2017, Lawson and Lücker 2018) which may allow for their co-existence. In fact, in three ND systems (i.e., DCWM, NEU and YAN) and the IFAS ND system (i.e., JAMMSM) the concentrations of AOB and comammox were positively correlated (R = 0.53 (DCWM), 0.41 (NEU), 0.91 (YAN), 0.60 (JAMMSM), p<0.01), suggesting the potential for some level of cooperation between the two. Further, like other lineage II *Nitrospira*, comammox bacteria demonstrate potential to exhibit diverse metabolic capacities (Daims et al. 2016, Lawson and Lücker 2018) including genes involved in urea uptake and urease for conversion of urea to ammonia (Daims et al. 2016, Koch et al. 2015, Lawson and Lücker 2018). Thus, it is plausible that the combination of greater metabolic diversity (e.g., ability to utilize urea) and higher affinities for ammonia of comammox bacteria compared to that of AOB, may ensure that comammox bacteria may have preferential access to ammonia (or other electron donors) made available via increased biomass decay products at longer SRTs. This could likely explain the increase in the abundance of comammox bacteria and not AOB or NOB with increasing SRT.

The second insight is that systems with comammox bacteria may have higher ammonia removal efficiencies compared to nitrogen removal systems where they are absent. It is important to note that this statistically significant observation is confounded by the fact that systems with comammox bacteria also exhibited higher SRT. Thus, it is difficult to disentangle the role of comammox bacteria from that of SRT alone in explaining the differences in ammonia removal efficiency. Finally, we did not find any other significant correlations between comammox bacterial concentrations and proportions (of total bacteria and ammonia oxidizers) with any of the other measured process parameters except including loading rates, pH, temperature, and dissolved oxygen (DO) concentrations. The lack of association with DO concentrations is in contrast with previous studies (Beach and Noguera 2019, Roots et al. 2019). In fact, comammox bacteria were present and their abundances stable in systems with DO less than 1 mg/l (i.e., YAN, NEU) as well as those with DO levels significantly in excess of 2 mg/l (i.e., GRE, JAMM, WIL, KIN). It is important to note that correlations of comammox bacterial abundance and proportions with process parameters were estimated on a system-by-system basis, with a maximum of 12 data points per system. Thus, it is feasible that our correlation efforts were limited by the richness of our dataset on an individual system basis, where multiple factors may together play a role in influencing microbial community composition. Thus, while this does not eliminate the possibility that low DO levels (or other process parameters) impact comammox bacteria, it suggests that SRT plays a more prominent role compared to other process parameters in the selection of comammox bacteria in full-scale mainstream nitrogen removal systems.

## 4.0 Conclusions

- Comammox bacteria were not prevalent in side-stream PNA systems, including single stage systems that contain attached growth and suspended phase components. This is likely to be associated with the high ambient ammonia concentrations in PNA systems.
- All comammox bacteria detected in full-scale systems belonged to clade A comammox bacteria and are closely associated with *Ca* Nitrospira nitrosa. This study provides a novel primer set and qPCR assay targeting the *amoB* gene of clade A comammox bacteria.
- Comammox bacteria were prevalent in full-scale mainstream nitrogen removal systems with long SRT’s and/or systems with attached growth components. This finding is consistent with estimates of slower growth rates for comammox bacteria compared to canonical AOB and NOB.
- Increases in comammox bacterial abundance in systems with sufficient SRT and/or attached growth phase were not associated with a concomitant decrease in the abundance of canonical AOB or NOB indicating that they may occupy niche independent from that of canonical nitrifiers within complex nitrifying communities.
- We found no significant associations between DO concentrations and comammox presence/absence or concentration in this study. While this does not eliminate the possibility that low DO levels favor comammox bacteria in wastewater treatment systems, it suggests that SRT is the key variable driving the prevalence of comammox bacteria.

## 5.0 Acknowledgements

This research was supported by the Water Environment Reuse Foundation (Grant UR416) and The National Science Foundation (Award number: 1703089). The authors also acknowledge the operational personnel for assistance with sampling and process data sharing. The authors would also like to thank Prof. Kartik Chandran (Columbia University) and Prof. Holger Daims (University of Vienna) and Dr. Petra Pjevac (University of Vienna) for providing DNA extracts of *Nitrosomonas europaea* and *Ca* Nitrospira inopinata, respectively, for preparation for qPCR standards.

