## Supplemental Figures for "Long solids retention times and attached growth phase favor prevalence of comammox bacteria in nitrogen removal systems"

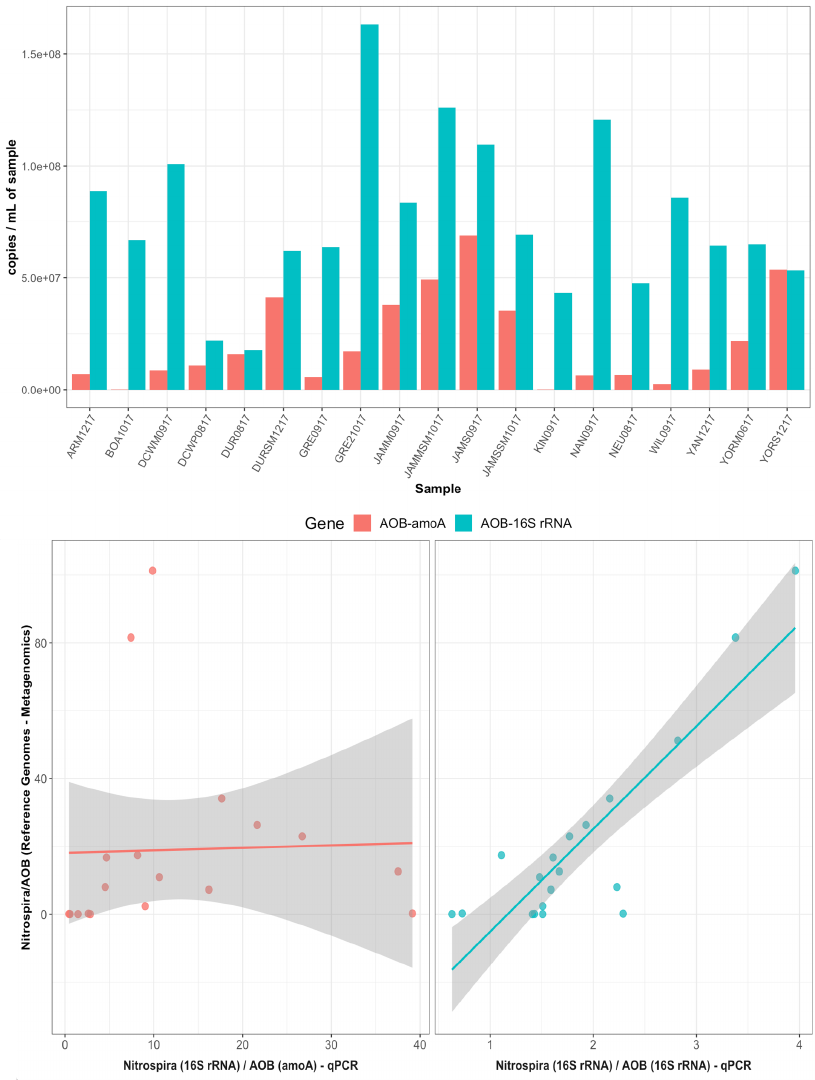

Figure S1: qPCR assays targeting the amoA gene did not concur with estimates from qPCR assay targeting the 16S rRNA gene of AOB (top panel). Further, ratios of Nitrospira to AOB (using amoA qPCR) estimates did not correlate with metagenomic analyses (bottom panel - left), while Nitrospira to AOB ratios using qPCR assays targeting the 16S rRNA gene were highly correlated with metagenomic data.
